# Organizing principles of pulvino-cortical connectivity in humans

**DOI:** 10.1101/205039

**Authors:** Michael J. Arcaro, Mark A. Pinsk, Janice Chen, Sabine Kastner

## Abstract

The pulvinar regulates information transmission to cortex and communication between cortical areas. The way the pulvinar interacts with cortex is governed by its intrinsic organization. Here, we show using fMRI that the human pulvinar is functionally heterogeneous, broadly separated into dorsal and ventral subdivisions based on characterization of response properties and functional connectivity with cortex. These differences mirrored the organization of the dorsal and ventral streams of visual cortex. The ventral subdivision of the pulvinar was functionally coupled with occipital and temporal cortex. The dorsal subdivision of the pulvinar was functionally coupled with frontal and parietal cortex. The dorsal subdivision was also coupled with the human-specific tool network and to the default mode network. The spatial organization of pulvino-cortical coupling reflected both the functional similarities and anatomical distances between cortical areas. Together, the human pulvinar appears to represent the entire visual system and the principles that govern its organization, though in a spatially compressed form.

**Author Contributions:** MA, MP, and JC collected data; MA and JC analyzed the data; MA, MP, JC, and SK wrote the paper.

## Introduction

The pulvinar is anatomically heterogeneous and extensively interconnected with visual cortex. As a general principle, cortical areas that are directly connected are also indirectly interconnected via the pulvinar (Shipp, 2003). Through this connectivity, it is thought that the pulvinar regulates corticocortical communication (Jones, 2001; Shipp, 2003; Saalmann et al., 2012). The function of the pulvinar’s influence on cortex is governed by its organization. Most of our understanding about the pulvinar comes from studies in non-human primates, though the broad organization of the human pulvinar appears to be similar to other primate species (Fig. 1). Across primates, the ventral pulvinar contains two well-defined maps of visual space (Allman et al., 1972; Bender, 1981; Li et al., 2013; Arcaro and Kastner, 2015). The ventral pulvinar is mainly connected with occipital visual areas and the dorsal pulvinar is connected with parietal and frontal regions(Shipp, 2003; Kaas and Lyon, 2007; Schmahmann and Pandya, 2008). Taken together, these data suggest a general distinction between the dorsal and ventral pulvinar.

The organizational principles governing pulvino-cortical connectivity may be guided by several factors. Within the monkey pulvinar, anatomical cortical connections appear to be topographically organized with neighboring parts of cortex projecting to neighboring parts of the pulvinar (Baizer et al., 1993). In the dorsal pulvinar, posterior parietal areas project to lateral portions of the dorsal pulvinar with anterior parietal areas projecting to more medial portions of the dorsal pulvinar (Fig. 1b)(Schmahmann and Pandya, 2008). In the ventral pulvinar, occipital cortex projects to anterior portions of the ventral pulvinar with temporal cortex projecting to posterior portions of the pulvinar (Fig. 1b)(Shipp, 2003). A recent study from our lab demonstrated a similar broad differentiation of dorsal and ventral pulvinar connectivity in humans using anatomical and functional imaging(Arcaro et al., 2015). How do these broad anatomical connectivity patterns relate to the functional organization of visual cortex? Based on prior anatomical studies, it might be expected that pulvino-cortical functional coupling purely reflects cortical distance with neighboring cortical areas projecting to neighboring parts of the pulvinar. For example, V1 may be connected with a region of the pulvinar close to V2’s connection, but at a further distance from V4’s connection. This could reflect the observation that neighboring cortical areas are frequently connected(Barbas and Pandya, 1989; Young, 1992). However, some cortical connections extend across large distances(Barbas and Mesulam, 1981; Markov et al., 2013). Pulvino-cortical connectivity may also reflect functional specialization of cortex with regions subserving similar functions projecting to the same parts of the pulvinar, regardless of cortical distance. For example, regions in parietal and frontal cortex involved in attentional control may overlap in their projections to the pulvinar. Such an organization is predicted based on recent physiological studies in monkeys showing that the pulvinar plays a role in synchronizing activity between cortical regions involved in a visual task(Saalmann et al., 2012). It is possible that these organizing principles coexist within the pulvinar and even vary as a function of subregion within the pulvinar.

Using neuroimaging, we investigated the functional organization of the pulvinar and its interactions with cortex in humans. We found that functional response properties across a variety of tasks and functional connectivity patterns (i.e., correlated fMRI signal) differed greatly between the ventral and dorsal pulvinar. Within the dorsal pulvinar, functional connectivity generally reflected the functional similarity of fronto-parietal areas, though some degree of cortical distance was also apparent. In contrast, within the ventral pulvinar, functional connectivity reflected cortical distance between occipital and temporal areas, and even differentiated specialized regions subserving similar functions (e.g., face patches in IT). These data provide a functional account of the principles guiding the pulvinar’s organization and pulvino-cortical functional coupling, revealing influences of both cortical distance and functional similarity. Our results suggest that the principles guiding pulvinar organization differ between dorsal and ventral subdivisions, and have important implications for the role of the pulvinar in facilitating cortical-cortical communication.

**Figure 1.**
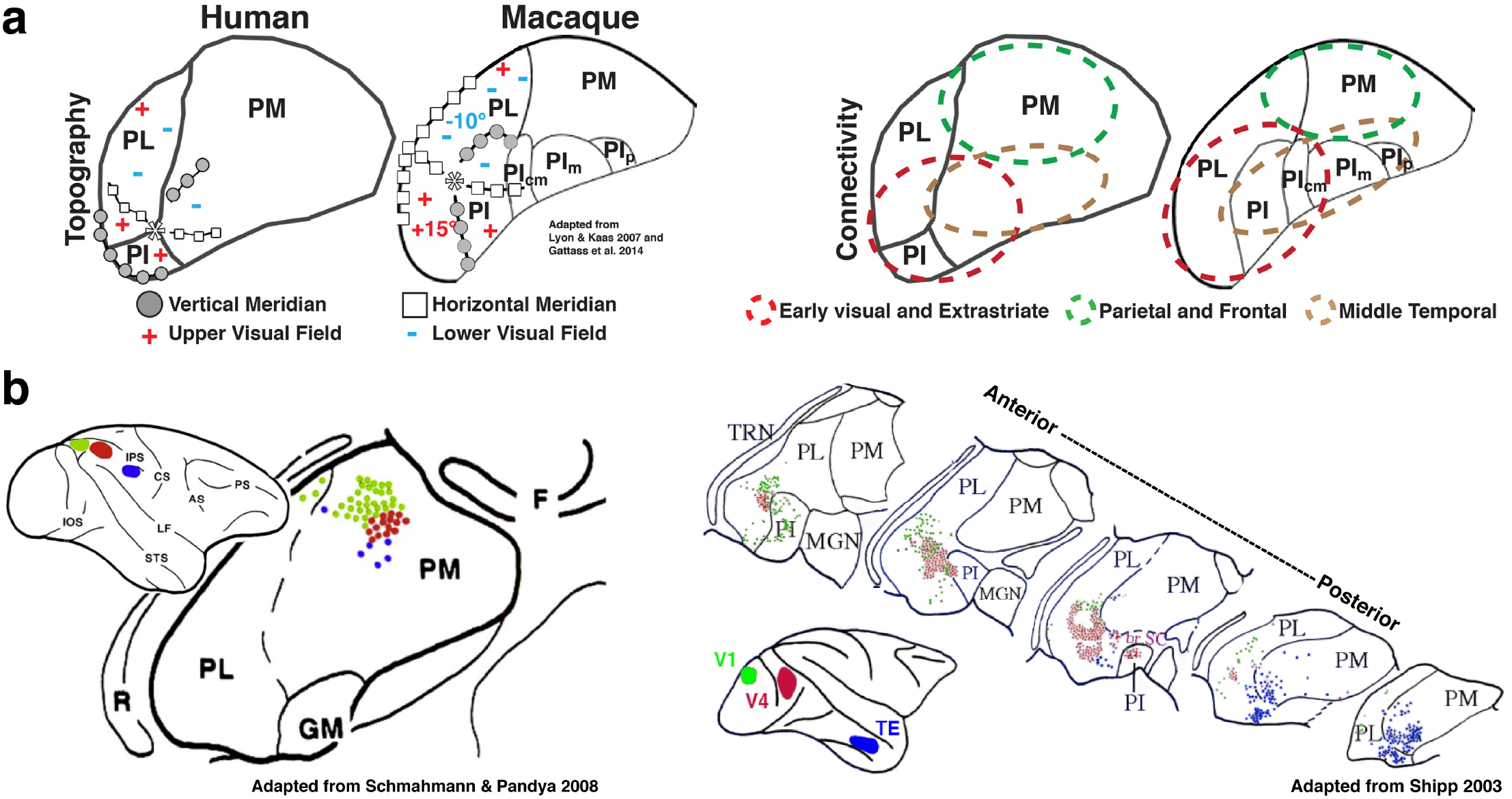
Organization of thalamo-cortical connections in humans and macaques. (a) Summary of retinotopic organization of the pulvinar (left) and broad connectivity with occipital, temporal, parietal, and frontal cortices (right).(b) Summary of parietal connectivity with dorsal pulvinar (left) and occipital-temporal connectivity with the ventral pulvinar (right). Connections are topographically organized with nearby cortical areas projecting to neighboring and partially overlapping portions of the pulvinar. (b, left) Image adapted from Schmahmann & Pandya, 2008. (b, right) Composite image adapted from Shipp 2003. To create this image, the V1 labels were extracted from Fig. 4a (Shipp 2003) and visually aligned to the V4 and TE labels from 4c based on the general shape of the pulvinar and the locations of lateral, inferior, and medial subdivisions. Specifically, the 5 posterior-most slices in Fig. 4a (Shipp 2003) were manually aligned to the 5 slices in the bottom of Fig. 4c. V1 labels were originally reported in Lysakowski et al. 1998 and V4 / TE labels were reported in Baleydier & Morel 1992.

## Materials and Methods

### Participants

Fifteen subjects (aged 20 – 36 years, seven females) participated in the study, which was approved by the Institutional Review Board of Princeton University. All subjects were in good health with no history of psychiatric or neurological disorders and gave their informed written consent. Subjects had normal or corrected-to-normal visual acuity. Thirteen subjects participated in task-free resting-state. In addition, all subjects participated in four experiments to localize cortical visual areas: (i) polar angle cortical mapping; (ii) eccentricity cortical mapping; (iii) memory-guided saccade mapping; (iv) object category localizer. Eleven subjects participated in a movie viewing experiment. Five subjects participated in the laterality and attentional modulation experiments. To localize subcortical visual field maps, nine subjects participated in a polar angle mapping session and five subjects participated in an eccentricity mapping session using scanning protocols optimized for subcortical structures (for more details, see Arcaro et al. 2015).

### Visual Display

The stimuli were generated on Macintosh G4 and G5 computers (Apple) using MATLAB software (MathWorks) and Psychophysics Toolbox functions (Brainard, 1997; Pelli, 1997). Stimuli were projected from either a Christie LX650 liquid crystal display projector (Christie Digital Systems) or a Hyperion PST-100984 digital light processing projector (Psychology Software Tools) onto a translucent screen located at the end of the Siemens 3T Allegra and Skyra scanner bores, respectively. Subjects viewed the screen at a total path length of ~60 cm through a mirror attached to the head coil. The screen subtended either 30°× 30° (Allegra), or 51°× 30° (Skyra) of visual angle. A trigger pulse from the scanner synchronized the onset of stimulus presentation to the beginning of the image acquisition.

### Experiments

*Resting State*. Thirteen subjects participated in two versions of resting state scans: (1) fixation; and (2) eyes closed. During the fixation scans, subjects were instructed to maintain fixation on a centrally presented dot (0.3 **°** diameter) overlaid on a mean grey luminance screen background for 10 minutes. During the eyes closed scans, the projector was turned off and subjects were instructed to keep their eyes closed for 10 minutes. Two runs were collected per resting state condition.

*Movie viewing*. A timescale localizer was used to delineate regions of the pulvinar that were sensitive to short-and long-timescales following established procedures(Hasson et al., 2008; Lerner et al., 2011; Honey et al., 2012; Chen et al., 2016). Eleven subjects viewed an audiovisual movie clip from the 1975 commercial film *Dog Day Afternoon*(Lumet, 1975). Subjects were instructed to attend to the movie and to freely view it. Movie stimuli subtended 20**°** horizontally and 16**°** vertically. A 5 min. 45 s clip of the film was presented as well as a temporally scrambled version of the stimulus where the clip was broken into segments spanning 0.5 – 1.6s and reordered. See(Chen et al., 2016) for more details.

*Laterality and attentional modulation*. Five subjects participated in an experiment designed to measure contralateral tuning and attentional modulation. Flickering checkerboard stimuli were presented to either the right or left visual hemifield. Stimuli subtended 15**°** horizontally and vertically. Subjects were instructed either to attend to a central fixation point (0.5**°**) and to detect changes in its luminance, or to maintain central fixation while covertly attending to one of the hemifields and to detect changes in luminance within a (varying) focal region of the checkerboard. The subjects’ task was varied between blocks. Blocks lasted 16s and were interleaved with 16s periods where subjects maintained central fixation on a blank grey screen.

### Functional Localizers

*Retinotopic Mapping*. All subjects participated in a single scan session in which polar angle and eccentricity representations were measured across cortex using a standard traveling wave paradigm consisting of a wedge or annulus, respectively (Swisher et al., 2007; Arcaro et al., 2009; Arcaro et al., 2011; Wang et al., 2014). A subset of these subjects participated in two additional scan sessions in which polar angle and eccentricity representations were measured within the pulvinar using a similar paradigm, but scanning protocols optimized for subcortical structures (Arcaro, 2015). Stimuli mapped the central 15° of the visual field. Due to limitations of the scanner bore size and viewing angle, peripheral representations beyond 15° were not mapped nor included in any analyses. Each run consisted of eight 40s cycles. For each subject, 2-5 runs were collected for cortical mapping and 8-10 runs were collected for pulvinar mapping. Fourier analysis (Bandettini et al., 1993; Engel et al., 1997; Sereno et al., 2001) was used to identify voxels that were sensitive to the spatial position (i.e., polar angle) of a peripheral cue during the task. Early visual and extrastriate areas V1, V2, V3, hV4, V3A–B, VO1–2, PHC1-2, LO1-2, TO1-2 were defined using standard criteria reported previously (Sereno et al., 1995; DeYoe et al., 1996; Engel et al., 1997; Brewer et al., 2005; Wandell et al., 2007; Arcaro et al., 2009). Pulvinar visual field maps, vPul1 and vPul2, and other subcortical visual field maps were defined using standard criteria previously published (Schneider et al., 2004; Schneider and Kastner, 2005; Arcaro, 2015). See (Arcaro et al., 2009) and (Arcaro et al., 2015) for details on cortical and pulvinar retinotopic mapping, respectively.

*Memory-guided saccade task*. All subjects participated in a single scan session in which a memory-guided saccade task was used to localize topographically organized areas in parietal and frontal cortex (Kastner et al., 2007; Konen and Kastner, 2008a, b). This task incorporates covert shifts of attention, spatial working memory, and saccadic eye movements in a traveling wave paradigm. The detailed description of the design and scanning parameters is provided in (Kastner et al., 2007; Konen and Kastner, 2008a). Briefly, subjects had to remember and attend to the location of a peripheral cue over a delay period while maintaining central fixation. After the delay period, subjects had to execute a saccade to the remembered location and then immediately back to central fixation. The target cue was systematically moved on subsequent trials either clockwise or counterclockwise among eight equally spaced locations. Each run was composed of eight 40 s cycles of the eight target position sequence. A total of eight runs were collected in a single scan session for each subject. Fourier analysis (Bandettini et al., 1993; Engel et al., 1997; Sereno et al., 2001) was used to identify voxels that were sensitive to the spatial position (i.e., polar angle) of a peripheral cue during the task. Parietal and frontal areas IPS0-5, SPL1, FEF, and IFS were defined using criteria previously published (Kastner et al., 2007; Swisher et al., 2007; Konen and Kastner, 2008b).

*Object localizer*. All subjects participated in a single scan session in which a standard object category localizer was used to define the occipital face area (OFA; (Puce et al., 1996)), fusiform face area (FFA; (Kanwisher et al., 1997; McCarthy et al., 1997)), anterior temporal face-selective area (AT;(Pinsk et al., 2009)); extrastriate body area (EBA; (Downing et al., 2001)), fusiform body area (FBA; (Peelen and Downing, 2005)), and parahippocampal place area (PPA; (Aguirre et al., 1996; Epstein and Kanwisher, 1998)). Briefly, grayscale pictures of objects (~12 × 12) from different categories were presented in 15s blocks, each containing 20 stimuli (350ms duration, 400ms interstimulus interval). Subjects viewed 12 blocks per stimulus category over the course of 4 runs. During stimulus presentation, subjects maintained central fixation and performed a one-back task indicating the repeated presentation of an object. Stimuli for each block were drawn from one of five categories: faces, houses, headless bodies, intact generic objects, and scrambled pictures of generic objects. The OFA was defined as a region within the occipitotemporal sulcus that showed significantly stronger activity during the presentation of faces compared with intact object stimuli (*p* < 0.0001). The FFA was defined as a region within the lateral fusiform sulcus based on the same statistical criteria. For many subjects, this region included two distinct sub-regions in close anatomical proximity (FFA-1/2, (Pinsk et al., 2009); pFus/mFus (Grill-Spector and Weiner, 2014)). AT was defined as a region within anterior temporal cortex based on the same contrast, though with a slightly lower threshold (*p* < 0.01). The EBA was defined as a region within the lateral occipitotemporal cortex that showed significantly stronger activity during the presentation of headless bodies compared with intact object stimuli (*p* < 0.0001). The EBA partially overlapped retinotopic areas LO2 and TO1. The FBA was defined as a region within the fusiform sulcus based on the same statistical criteria. In several subjects, the FBA partially overlapped the FFA. Overlapping voxels were assigned to either the FFA or FBA based on contrasting activity during face and headless body presentations. The PPA was defined as a region within the posterior parahippocampal cortex within the collateral sulcus and along the medial fusiform sulcus that showed significantly stronger activity during the presentation of houses compared with intact object stimuli (*p* < 0.0001). The PPA largely overlapped with retinotopic areas PHC1 -2. Since the PPA was localized using a different experiment and may represent functional dissociations within this part of cortex, voxels were not restricted to non-overlapping portions. The TOS partially overlapped with retinotopic areas V3B, IPS0, and LO1. Since the areas EBA, PPA, and TOS were localized using a different experiment and may represent functional dissociations, voxels were not restricted to portions of cortex non-overlapping with retinotopic areas. Group-level regions were identified using a mixed effects meta-analysis (AFNI’s 3dMEMA).

*Additional areas used in pulvino-cortical correlation analyses*. The Posterior Cingulate/Precuneus (PreC) and inferior parietal areas (IPL) were identified based on relative anatomical location and MNI coordinates from prior studies on the default mode network(Buckner et al., 2008). The lateral temporal (latTemp) and anterior parietal (antPPC) areas were identified based on relative anatomical location and MNI coordinates from prior studies on the tool network(Mruczek et al., 2013). Lateral (latPPC) and medial parietal (medPPC) areas were identified as regions located cortically in-between the IPS2-4 and the IPL and PreC, respectively, that are functionally dissimilar. MNI coordinates were projected onto each subject’s native space anatomical surface. An ROI spanning 6mm on the surface around each MNI- or anatomically-defined region was drawn for each area. Areas of overlap with surrounding retinotopic maps were excluded. For overlap between latPPC and IPL (medPPC and PreC), the border was adjusted to the midpoint. ROI size did not greatly affect analyses and results were comparable for larger (8mm) and smaller (4mm) ROI sizes.

*HCP multi-modal parcellation*. An atlas of 180 areas across the cortical surface of each hemisphere(Glasser et al., 2016) in MNI space was aligned to each subject’s native anatomical image using non-linear anatomical registration procedures outlined below. These 180 areas were used in a secondary analysis to evaluate pulvinar functional connectivity patterns based of correlations across the whole cortical surface.

### Data acquisition

Functional MR images were acquired with a gradient echo, echo planar imaging (EPI) sequence using an interleaved acquisition. The specific parameters for each scan session are outlined below.

For the laterality / attentional modulation scans, data were acquired with a Siemens 3T Allegra scanner using a circularly polarized head coil (Siemens, Erlangen, Germany). Laterality and attentional modulation experiments, 25 oblique slices were acquired using an EPI sequence with a 128-square matrix (slice thickness 2.5mm, interleaved acquisition) leading to an in-plane resolution of 1.5 × 1.5 mm^2^ [field of view (FOV) = 256 × 256 mm^2^; repetition time (TR) = 2.0s; echo time (TE) = 40 ms; flip angle (FA) = 90**°**].

For all other experiments, data were acquired with a Siemens 3T Skyra scanner using 20-channel phased-array head(16)/neck(4) coil (Siemens, Erlangen, Germany). All functional acquisitions used a gradient echo, echo planar sequence with a 64-square matrix (slice thickness of 4mm, interleaved acquisition) leading to an in-plane resolution of 3 × 3 mm^2^ [FOV = 192 × 192 mm^2^; GRAPPA iPAT = 2; 32 slices per volume for resting state and 27 for movie stimuli; TR = 1.8 s for resting state and 1.5 s for movie scans; TE = 30 ms; FA = 72**°**]. High-resolution structural scans were acquired in each scan session for registration to surface anatomical images (MPRAGE sequence; 256-square matrix; 240 × 240 mm^2^ FOV; TR, 1.9 s; TE 2.1 ms; flip angle 9**°**, 0.9375 × 0.9375 × 0.9375 mm^3^ resolution).

### Data analysis

*Preprocessing*. Data were analyzed using AFNI (Analysis of Functional NeuroImages, RRID:nif-0000-00259; Cox, 1996), SUMA (Saad and Reynolds, 2012), FSL (FSL, RRID:birnlex_2067; Smith et al., 2004; Woolrich et al., 2009; Jenkinson et al., 2012; http://fsl.fmrib.ox.ac.uk/fsl/fslwiki/), FreeSurfer (FreeSurfer, RRID:nif-0000-00304; Dale et al., 1999; Fischl et al., 1999; http://surfer.nmr.mgh.harvard.edu/), and MATLAB (MATLAB, RRID:nlx_153890). Functional data were slice-time and motion corrected. Motion distance (estimated by AFNI’s 3dvolreg) did not exceed 1.0 mm (relative to starting head position) in any of the 6 motion parameter estimates (3 translation and 3 rotation) during any run for any subject.

*Laterality and attentional modulation experiment*. Data were spatially filtered with a 4mm (FWHM) Gaussian kernel, which increased SNR while maintaining good anatomical localization of signals within the pulvinar (Figure 1). A multiple regression analysis (AFNI’s 3dDeconvolve) in the framework of a general linear model was performed. Each stimulus condition was modeled with square-wave functions matching the time course of the experimental design convolved with a hemodynamic response function. Additional regressors that accounted for variance due to baseline shifts between time series, linear drifts, and head motion parameter estimates were also included in the model. Brain regions that responded more strongly to right or left visual fields were identified by contrasting blocks of checkerboard stimulation to the right or left visual field while the subject maintained a central fixation. Brain regions that responded more strongly during covert attention were identified by contrasting blocks of covert attention to the checkerboard stimuli vs. blocks of central fixation during visual stimulation. In each hemisphere, covert attention contrasts were only considered for attention to the contralateral visual field. Contralateral tuning was assessed by computing a d prime index (d’), defined by the following formula

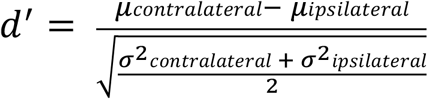

where *μcontralateral* and *μipsilateral* are the average responses to contralateral visual stimuli (during central fixation); *σcontralateral* and *σipsilateral* are the SDs. Attentional modulation was calculated using the same formula between covert hemifield attention and central fixation conditions.

*Additional preprocessing for correlation analyses*. Several additional steps were performed on the data: (1) removal of signal deviation > 2.5 SDs from the mean (AFNI’s 3dDespike); (2) temporal filtering retaining frequencies in the 0.01-0.1 Hz band; (3) linear and quadratic detrending; and (4) removal by linear regression of several sources of variance: (i) the six motion parameter estimates (3 translation and 3 rotation) and their temporal derivatives, (ii) the signal from a ventricular region, and (iii) the signal from a white matter region. To avoid partial volume effects with surrounding grey matter, ventricular and white matter regions were identified by hand on each subject’s mean EPI image. These are standard preprocessing steps for resting-state correlation analyses (Vincent et al., 2007; Yeo et al., 2011), though our results were not dependent on these preprocessing steps, and correlation analyses on the raw data yielded qualitatively and statistically similar results. Global mean signal (GMS) removal was not included in the analysis reported here given concerns about negative correlations (Fox et al., 2009; Murphy et al., 2009; Saad et al., 2012), though inclusion of GMS removal yielded statistically similar results. To minimize the effect of any evoked response due to the scanner onset, the initial 21.6 s were removed from each rest scan. To extract the mean signal from each cortical area, voxels that fell between the gray and white matter boundaries were mapped to surface model units (nodes). The mean signal from each cortical area was extracted into MATLAB for correlation analyses. Data within the thalamus was then spatially filtered with a 4mm (FWHM) Gaussian kernel, which increased SNR and improved correspondence of correlation patterns between subjects while maintaining good anatomical localization of signals within the pulvinar.

*Inter-subject correlations*. For the movie viewing experiment, an inter-subject correlation (ISC) analysis approach was used (Hasson et al. 2004; Hasson et al., 2008; Chen et al., 2016). This analysis provides a measure of the consistency of a response to the temporally complex stimulus (i.e., naturalistic audiovisual movie) by comparing the BOLD response across different subjects. ISC helps avoid the contribution of idiosyncratic responses to correlation patterns and circumvents the need to specify a model for the neuronal processes in any given brain region during movie watching. For ISC, repetitions for each condition were averaged within subject. Data were then transformed to MNI space and voxel-wise correlations were computed between the responses in each subject with the average of all other subjects for each condition separately. This yielded a voxel-wise measure of the consistency of brain activity evoked by the audiovisual movie clip. Group average correlation maps were calculated and the difference was computed between intact and scrambled correlation maps. See(Chen et al., 2016) for more details.

*Pulvino-cortical functional connectivity analyses*. We applied a two-step Pearson correlation analysis to identify cortical areas whose spatial profile of correlations with all other cortical areas (referred to as cortical area correlation profile) was similar to the cortical area correlation profile of individual voxels in the pulvinar. First, *temporal* correlations were performed between each cortical area and between cortical areas and the pulvinar. To identify the cortical correlation profile of individual cortical areas, the mean timeseries of each cortical area was correlated with the mean timeseries of every other cortical area in each subject. To identify the cortical correlation profile of the pulvinar in each subject, the mean timeseries of each cortical area was correlated with the timeseries of each voxel within a pulvinar mask.

Next, we compared the *pattern* of cortical correlations (profiles) for individual pulvinar voxels with those of individual cortical areas. For each subject, the cortical correlation profile of each pulvinar voxel was correlated to the pseudo-group average cortical correlation profile of each cortical area, where the average excluded that subject. This yielded a measurement of similarity between each cortical area’s cortical correlation profile and every pulvinar voxel’s cortical correlation profile, which we refer to as the *pulvino-cortical connectivity*. Significant positive similarity in the pulvino-cortical functional connectivity was taken as evidence for connectivity between a pulvinar voxel and a given cortical area. Individual subject anatomical volumes were then aligned using a two-step linear (AFNI’s 3dAllineate), nonlinear (AFNI’s 3dQwarp) registration procedure, and each subject’s pulvino-cortical functional connectivity profile was transformed into MNI space. Voxel - wise, one sample t-tests were used to assess statistical significance. See (Arcaro et al., 2015) for additional details on this approach. Even for cortical areas that were investigated previously(Arcaro et al., 2015), the computation of each pulvino-cortical connectivity map differs in this study due to the inclusion of several additional parietal, temporal, and frontal areas, which changes the size of the cortical connectivity profile. Though the cortical connectivity profile has been greatly expanded in this study, the resulting pulvino-cortical connectivity maps for areas previously reported(Arcaro et al., 2015) were very similar, demonstrating the robustness of these connectivity measures.

*HCP connectivity profile*. As a control analysis to ensure that the connectivity patterns for the 39 areas used in the main analysis were not dependent on the particular ROIs, areal connectivity profiles were calculated using an 180 ROI parcellation of the whole cortical surface(Glasser et al., 2016).

*Overlap*. To compare the spatial localization of correlation patterns within the pulvinar, the overlap of pulvino-cortical connectivity maps was computed for all areas pairs with Dice’s coefficient, *2*|*A*-*B*| / (|*A*|+|*B*|). The group average map for each cortical area was threshold at *p* < 0.05 and binarized. Degree of overlap was assessed relative to the entire anatomical volume of the pulvinar.

*Multidimensional scaling*. To assess the structure of the pulvino-cortical connectivity, correlations were calculated between the pulvino-cortical connectivity maps of each cortical area. This high-dimension 39 × 39 connectivity matrix was converted to a dissimilarity (Euclidean distance) matrix and classical multi-dimensional scaling (MDS) was applied. The first two principal dimensions were visualized (Fig. 2). To compare the structure of the first two principal dimensions between hemispheres and across subjects, procrustes analysis was performed. For hemispheres, the left hemisphere was transformed to the right hemisphere space. Individual subjects were transformed to the group average space. Permutation tests were performed for the individual subject transformations. For each subject, the area labels for the two principal dimensions were scrambled and procrustes analysis was performed on the permuted data. This permutation was computed for each subject 1000 times. To test whether the group data were representative of individual subjects, two versions of the permutation test were performed. (1) Labels were permuted across all areas to test whether the structure of the individual subjects was similar to the pseud-group structure (minus that subject) above chance. (2) Labels were permuted within each cluster from the group analysis to test whether the structure of individual subjects within each cluster was similar to the pseudo-group (minus that subject). The goodness of fit (SSE; summed square of residuals) to the pseudo-group data for each subject’s real data was compared with the goodness of fit for the permuted data.

*Clustering*. Clustering on the group average data was performed using a spectral (eigendecomposition) algorithm(Newman, 2006) from the Brain Connectivity Toolbox(Rubinov and Sporns, 2010). This algorithm automatically subdivides a (weighted) network into non-overlapping groups that maximizes the number of within-group links and minimizes the number of between group links. That is, the algorithm determines the optimal cluster size, which is an advantage over other clustering approaches, such as k-means. In practice, we found that k-means with a *k* matched to the cluster size returned from this spectral clustering approach yielded similar divisions that would not change interpretation of our results.

*Cortical distance*. Euclidean distances between the group average centroids for all 39 areas were calculated using AFNI’s SurfDist. Cortical distance measures were correlated with Dice’s coefficient (Sorensen index) and with the Euclidean distances between peaks in the pulvino-cortical connectivity maps.

### Experimental design and statistical analysis

Thirteen subjects participated in resting state scans. Individual and group-level analyses were performed. Patterns of co-fluctuations in the BOLD signal were assessed within subject using Pearson correlation. Group-level statistical significance was assessed for the spatial pattern of pulvinar correlations by Fisher transforming individual subject correlations and performing one-sample, two-tailed t-tests. Data were corrected for multiple comparisons (FDR) at p < 0.05. MDS and clustering analyses were performed on the individual and group-average data. Pearson correlations were performed between the spatial pattern of correlations and cortical distance. Eleven subjects participated in a movie viewing experiment. Patterns of co-fluctuations in the BOLD signal were assessed between subjects using a Pearson’s correlation. Five subjects participated in the contralateral tuning and attentional modulation experiment. Response magnitudes and significance were estimated for each subject using a regression analysis. Group-level statistical significance was assessed across subjects using a one-sample, two-tailed t-tests. Data were corrected for multiple comparisons (FDR) at p < 0.05. A d prime index was used to calculate degree of contralateral tuning and attentional modulation across subjects. Statistical significance was assessed across subjects using a two-sample, two-tailed t-tests. Fifteen subjects participated in the object localizer experiment. Response magnitudes and significance were estimated for each subject using a regression analysis. Group-level statistical significance was assessed across subjects using a mixed effects meta-analysis that models both within- and across-subject variability.

## Results

### Differential functional response profiles in dorsal and ventral pulvinar

Responses within the ventral and dorsal pulvinar varied as a function of spatial location, attentional allocation, and stimulus content (Fig. 2a). The ventral pulvinar responded robustly to visual stimulation (flickering checkerboard) and showed a greater response to contralateral (vs. ipsilateral) visual stimulation while subjects performed a dimming task at a central fixation point (Fig. 2a, left, *p* < 0.05, FDR corrected). The anatomical extent of these contralateral maps corresponded to the locations of the two visual field maps within the ventral pulvinar described previously (Arcaro et al., 2015; DeSimone et al., 2015). In contrast, the dorsal pulvinar responded weakly to visual stimulation and did not show a clear visual hemifield preference in individual subjects or the group average contrast maps. To further quantify the visual responses within the pulvinar, regions of interest within the ventral and dorsal pulvinar were identified based on group average retinotopic maps and functional connectivity with parietal and frontal cortex, respectively(Arcaro et al., 2015). The degree of contralateral tuning, as assessed by a d prime laterality index contrasting evoked activity from contralateral and ipsilateral visual stimulation (Fig. 2a, right), was much greater in the ventral pulvinar as compared with the dorsal pulvinar (t(4) = 5.96, *p* < 0.01, n = 5). Responses in the dorsal and ventral pulvinar also were differentiated based on the degree of attentional modulation. The dorsal pulvinar showed greater attentional modulation than the ventral pulvinar, as assessed by a d prime attentional modulation index contrasting contralateral visual stimulation during covert attention from contralateral stimulation during central fixation (t(4) = 3.16, *p* < 0.05, n = 5). These results are consistent with prior literature showing contralateral tuning and attentional modulation in the pulvinar(Smith et al., 2009). Together, these data show a differentiation between the ventral and dorsal pulvinar based on visual responsiveness and attentional modulation, mirroring the broad distinctions of occipital and fronto-parietal cortices.

The consistency of responses to an audiovisual input also varied between the dorsal and ventral pulvinar. Response consistency was assessed using an inter-subject correlation (ISC) approach(Hasson et al., 2008). Responses in both the dorsal and ventral pulvinar during viewing of intact naturalistic movies were consistent across individuals (Fig. 2b; mean r > 0.15, n = 11). However, only the ventral pulvinar showed consistent responses across individuals between viewings of scrambled versions of the same movie where the temporal structure and narrative flow was disrupted. Subtracting the response consistency of the scrambled movie from the intact movie yielded focal bilateral clusters of voxels within the dorsal pulvinar (r difference > 0.15, n = 11). These results mirror a previously established hierarchy of temporal receptive windows across occipital, parietal, and frontal cortex(Hasson et al., 2008; Honey et al., 2012). That is, activity in the ventral pulvinar appears to reflect moment-to-moment transitions in visual statistics, similar to occipital visual areas, whereas activity in the dorsal pulvinar reflects attentional and contextual information, similar to frontal and parietal areas.

Finally, activity in the dorsal and ventral pulvinar differentiated based on stimulus category selectivity. When comparing responses to viewing face and scene images, focal bilateral clusters of voxels in the posterior, ventral pulvinar showed a preference for faces vs. scenes (Fig. 2c, *p* < 0.05, FDR corrected, n = 15). The same region of the ventral pulvinar also showed a preference for faces vs. objects and faces + bodies vs. objects + scenes (*p* < 0.05, FDR corrected, n = 15). Voxels showing a preference for scenes vs. faces were identified lateral to the face/body sensitive clusters in both hemispheres. These data indicate a role for the posterior ventral pulvinar in high-level visual processing. Across all experiments, the functional response properties of the dorsal and ventral pulvinar broadly reflect functional distinctions between the ventral and dorsal visual streams.

**Figure 2.**
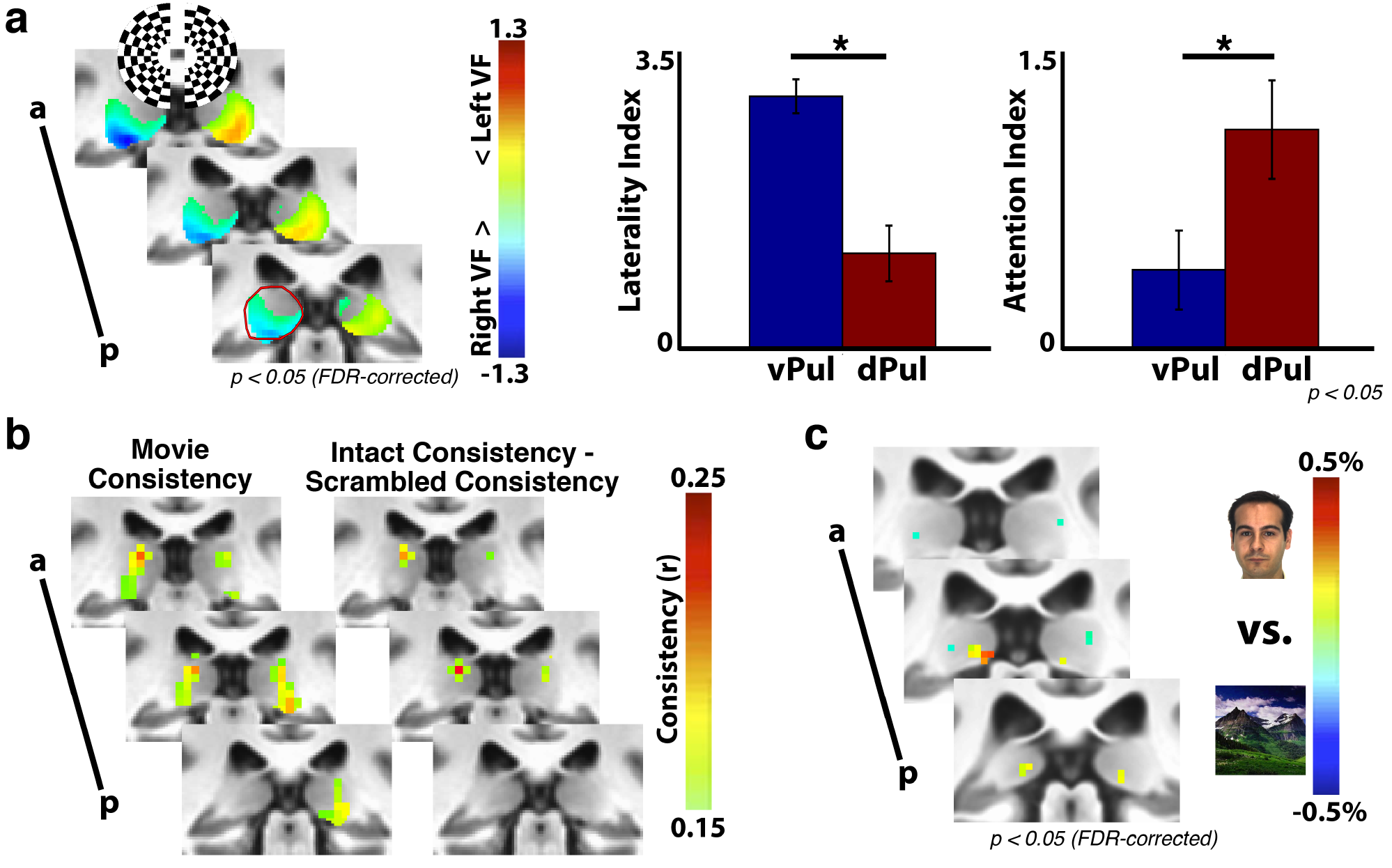
Functional distinctions within and between the dorsal and ventral pulvinar. (a) The ventral pulvinar represented contralateral visual space (left, p < 0.05, FDR corrected, n = 5), while the dorsal pulvinar showed greater attentional modulation (right, t(4) = 3.16, p < 0.05, n = 5). Anatomical extent of the left hemisphere pulvinar in posterior-most slice outlined in dark red. (b) Repeated presentations of intact movie stimuli evoked consistent responses in both the ventral and dorsal pulvinar. Only the dorsal pulvinar showed greater consistency to repeated presentations of intact vs. scrambled movies (r > 0.15, n = 11). (c) A posterior medial portion of the ventral pulvinar responded preferentially to face vs. scene stimuli (p < 0.05, FDR corrected, n = 15).

### Differential functional connectivity between the dorsal and ventral pulvinar

The differences in response selectivity between the dorsal and ventral pulvinar were reflected in their functional connectivity patterns with visual cortex. We assessed within-hemisphere correlated activity between the pulvinar and 39 cortical areas across occipital, temporal, parietal and frontal cortices using Pearson correlations. These areas tile visual cortex and were chosen as they share broad functional similarity (i.e., involved in various visual tasks) yet vary in their specializations and cortical location. Correlations for a subset of these areas were previously reported(Arcaro et al., 2015) using a much more limited dataset (See Materials and Methods: *Pulvino-cortical functional connectivity analyses*). For most cortical areas, the mean time-series was positively correlated with the time-series of many voxels in the pulvinar. In addition, most cortical areas were positively correlated with each other, making it difficult to evaluate the specificity of the correlated signal. We assumed that signals between directly connected areas should be more correlated than between indirect or weakly connected areas. Therefore, we reasoned that using the spatial correlation pattern across all areas should be a more sensitive measure (vs. pairwise temporal correlations). To do this, we first calculated the mean timeseries of each cortical area and computed temporal correlations between all cortical area pairs (referred to as cortical area correlation profile). We then computed the temporal correlation between the mean timeseries of each cortical area with the timeseries for each voxel in the pulvinar (voxel-wise pulvino-cortical temporal correlation profile). For each subject, we then compared the pattern of the voxel-wise pulvino-cortical temporal correlations with the (leaving that subject out) pseudo-group average cortical area correlation profile, yielding a voxel-wise measurement (spatial map) of the similarity between each cortical area’s correlation profile and every pulvinar voxel’s cortical area correlation profile. This measurement is referred to as the pulvino-cortical connectivity. A complete pulvino-cortical connectivity map within the pulvinar was generated for each cortical area in each subject and used for all subsequent analyses. [See (Arcaro et al., 2015) for more details.]

The group average pulvino-cortical connectivity (n = 13) broadly reflected the distinction between dorsal and ventral visual streams (Fig. 3). The similarity between each cortical area’s pulvino-cortical connectivity map was calculated, then averaged across hemispheres. Multidimensional scaling (MDS) illustrates the similarity of group average pulvino-cortical connectivity between cortical areas (Fig. 3a). Data were clustered using a spectral clustering algorithm(Newman, 2006; Rubinov and Sporns, 2010). The algorithm determined an optimal cluster size of 2. One cluster comprised all occipital and temporal areas (with the exception of a tool-selective area in anterior lateral temporal cortex). The other cluster comprised all parietal and frontal regions (with the exception of IPS0) as well as the retrosplenial cortex (RSC). For both clusters, most areas were linked with many other areas within the cluster (Fig. 3a) and had weak (r < 0.15) or no (r <= 0) links between clusters. The notable exception was RSC, which had several links with both clusters, indicating that it may serve as a bridge between dorsal and ventral subdivisions. Mean pulvino-cortical connectivity maps were calculated for both clusters (Fig. 3b).

To visualize any structure of this cluster organization in anatomical space, the difference of the cortico-pulvinar connectivity between these cluster maps was calculated for each subject and averaged. One-sample two-tailed t-tests were conducted to identify voxels that showed a consistent difference in cortico-pulvinar connectivity across subjects (*p* < 0.05 FDR corrected, Fig. 3b). The group average peak connectivity within the pulvinar for both clusters was symmetrical between hemispheres and extended along the anterior-posterior and lateral-medial axes. The occipital-temporal (blue) cluster was most strongly associated with the ventral pulvinar and the frontal-parietal (red) cluster was most strongly associated with the dorsal pulvinar, mirroring previous anatomical distinctions in non-human primates (Baleydier and Morel, 1992; Kaas and Lyon, 2007).

**Figure 3.**
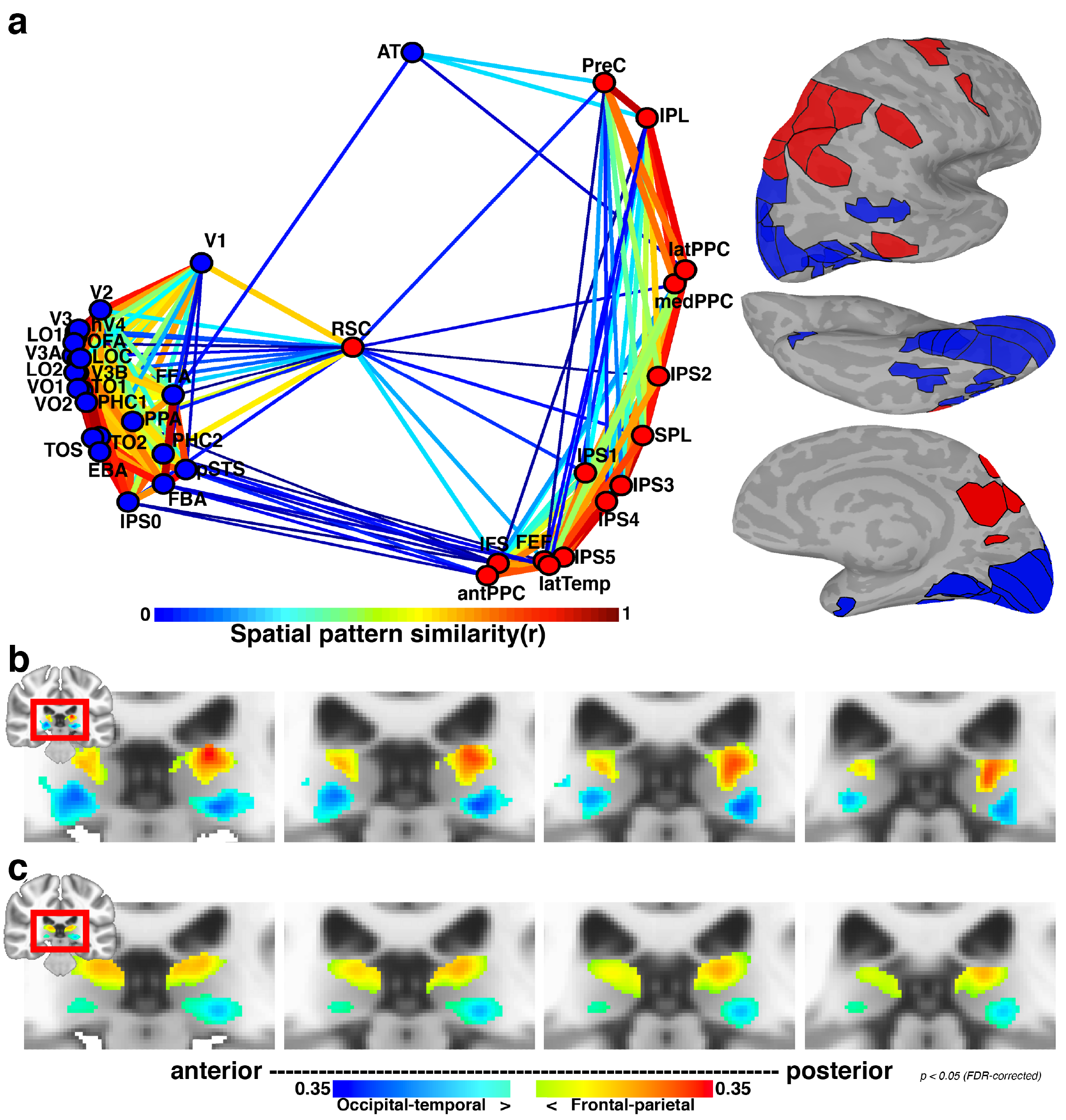
Organization of correlated activity between cortex and pulvinar. (a) 2D plot of multidimensional scaling on the group average (n=13) pattern similarity of pulvino-cortical connectivity between 39 cortical areas. Only ipsilateral correlations were considered. Data clustering yielded 2 groups. One cluster (blue) mainly contained occipital and temporal cortical areas associated with the ventral visual stream. The other cluster mainly contained frontal and parietal cortical areas associated with the dorsal visual stream. Line thickness and color coding reflects similarity strength between areas. Surface figures illustrate all cortical areas tested; fill color matched to assigned cluster. (b) The difference between the two cluster’s pulvino-cortical connectivity maps segmented the pulvinar into dorsal and ventral sections. Colors correspond to the blue and red clusters shown in (a), threshold at an p < 0.05 FDR corrected. Anatomical extent of the left hemisphere pulvinar outlined in dark red. (c) Same analysis as in (b), but from calculating correlations using the HCP 180 cortical areas as the cortical correlation profile.

The clustering of occipito-temporal and fronto-pareital areas within the ventral and dorsal pulvinar, respectively, held when evaluating functional connectivity patterns across all of cortex. Instead of computing cortical connectivity profiles based on correlations across the 39 areas tested, connectivity profiles were re-calculated for each of the 39 areas based on connectivity across a whole-brain segmentation of cortex into 180 areas (Glasser et al., 2016). Even when considering the connectivity profile across cortex, clustering of occipito-temporal areas and fronto-parietal areas remained (Fig. 3c). Notably, the clusters from the whole-brain connectivity profiles were less distinct (apparent by the smaller differences in the subtraction maps of Fig. 3c vs. 3b) and contained more cross cluster links than the clusters from the connectivity profiles restricted to visual areas. By inspection of the correlation matrices, this appeared to be due to the inclusion of several cortical areas in the connectivity profile that contained little to no correlations with areas in either the dorsal or ventral pulvinar cluster, thereby inflating the similarity of their connectivity profiles. This indicates that restricting the analysis to only visual areas was more sensitive at uncovering the connectivity structure of the pulvinar with visual cortex. Importantly, these broader clusters were also localized to the ventral and dorsal pulvinar, respectively (Fig. 3c), further demonstrating that this is a fundamental distinction within the human pulvinar.

The structure of pulvino-cortical connectivity was consistent across subjects (Fig. 4). The group average pulvino-cortical connectivity profile for each area was similar between hemispheres (mean r = 0.92 +/-0.02 across areas) and between averages from subsampling subjects (mean r = 0.73 +/-0.01 across areas from 500 iterations of split halves). MDS and clustering on the group average of each hemisphere separately yielded similar structure (Fig. 4a). Further, MDS on each subject’s pulvino-cortical connectivity yielded a more similar structure to the group average (Fig. 4b) than did random configurations from permutation testing (Methods: Multidimensional scaling). The goodness of fit (SSE) from procrustes analysis for each subject (mean sse = 0.04 +/0.005 s.e.m. across subjects; max = 0.08) was better than goodness of fit measures from permutation testing (min 99.99% across subjects = 0.62), indicating that the occipital/temporal and frontal/parietal distinction was present across individuals. Further, the goodness of fit for each subject was better than goodness of fit measures from permutation testing where the labels were permuted only within the clusters (min 99.99% across subjects = 0.11), indicating that the structure within each cluster was consistent across individuals.

**Figure 4.**
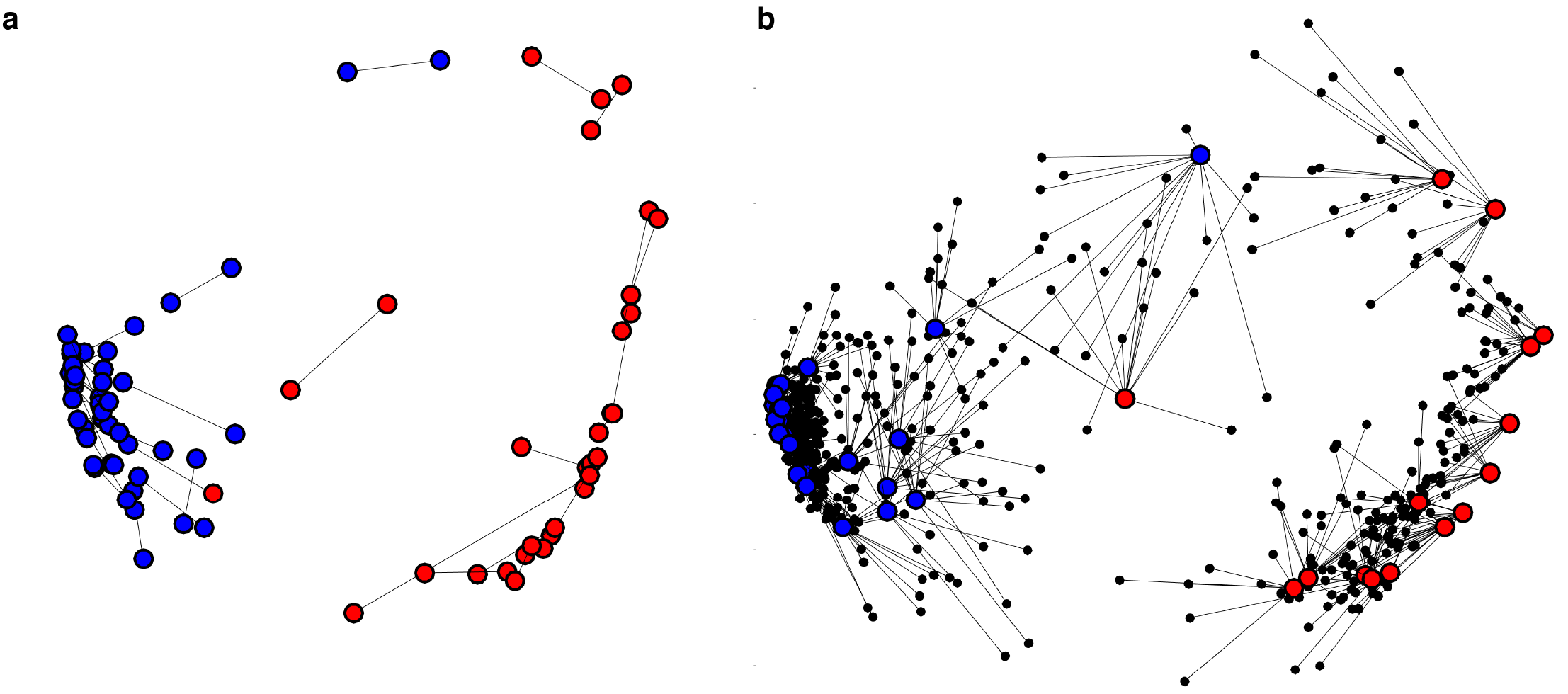
Consistency of pulvinar connectivity between hemispheres and across individuals. (a) Multidimensional scaling for left and right hemisphere pulvino-cortical functional connectivity. Procrustes analysis was performed to align MDS of the left hemisphere to the right hemisphere. Lines illustrate the distances between areas matched between hemispheres. Dots were color-coded based on clustering performed on each hemisphere separately. The only difference in clustering between hemispheres was area pSTS. (b) Multidimensional scaling for each subject’s pulvinar connectivity (averaged across hemisphere). Procrustes analysis was performed to align MDS of each subject to the group average. Lines illustrate distances of between individual subjects and the group average for matched areas. Dots were color-coded based on clustering performed on the group average.

The pulvino-cortical connectivity maps (threshold of *p* < 0.05 FDR-corrected from t-test across subjects) for neighboring cortical areas tended to have a good degree of overlap in the pulvinar and there was a linear relationship between the degree of overlap (Dice’s coefficient) and the cortical distance between seed areas (Fig. 5; r(739) = -0.48, p < 0.0001). This relationship was driven by correlations within the occipital-temporal cluster (r(274) = -0.63, p < 0.0001) and was not apparent within the frontal-parietal cluster (r(103)=-0.10, p=0.31). The difference between the two cluster correlation coefficients was significant (z = 5.52, p < 0.0001). Taken together, these data demonstrate a broad distinction between the dorsal and ventral pulvinar’s cortical coupling and the functional organization within each subdivision.

**Figure 5.**
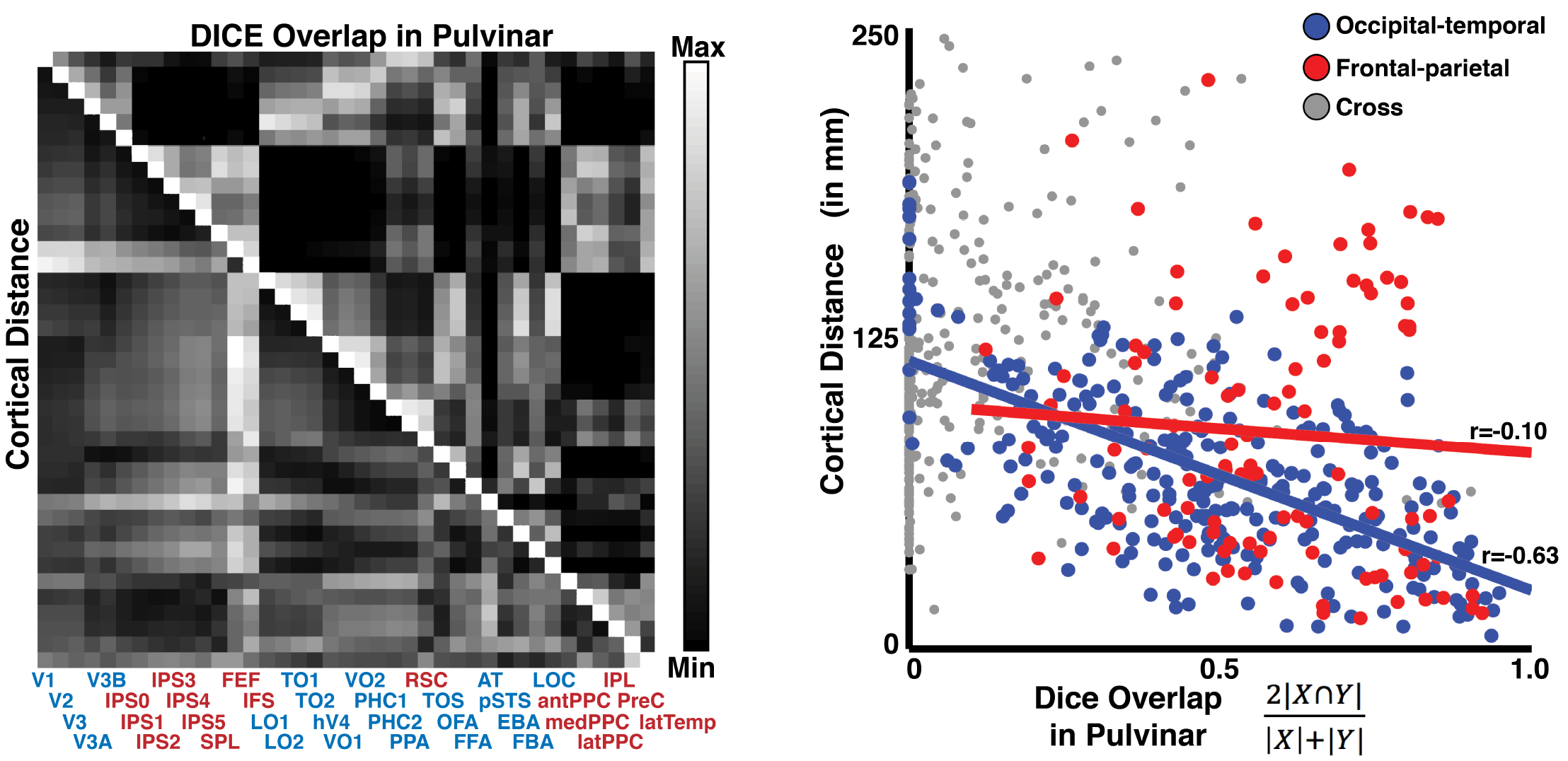
Relationship between cortical distance and cortical functional connectivity in the pulvinar. (left) Half matrices showing pairwise cortical distances and overlap in pulvino-cortical functional connectivity for all 39 cortical areas. Area labels are colored based on clustering in Figure 2. (right) Scatter plot of cortical distance vs. overlap of pulvino-cortical connectivity for area pairs that were both part of the ventral pulvinar cluster (blue), dorsal pulvinar cluster (red), or for area pairs between clusters (grey). The correlation between cortical distance and pulvino-cortical connectivity overlap was significant for areas within the occipital-temporal cluster (r = -0.63, p < 0.0001), but not within the frontal-parietal (r = -0.10, p > 0.10). For all area pairs, Dice’s coefficient, 2|A-B| / (|A|+|B|), was calculated.

### Organization of cortical connections within the dorsal pulvinar

Within the dorsal pulvinar, the organization of pulvino-cortical functional connectivity reflected the functional topography and response properties of parietal and frontal cortices. Consistent with prior work from our lab testing fewer areas (Arcaro et al., 2015), individual frontal and parietal areas were most strongly correlated with the dorsal pulvinar (Fig. 6a). Several individual pulvino-cortical connectivity maps greatly overlapped even between cortically distant areas such as IPS2 in parietal cortex and FEF in frontal cortex, while others appeared to be spatially distinct (IPL). To evaluate the finer-scale structure of these maps, the spatial locations of the peaks in pulvino-cortical functional connectivity maps were assessed for each area. Clustering on the distances between the peaks of pulvino-cortical connectivity maps revealed a finer structure of 3 clusters within the dorsal pulvinar (Fig. *6b*). The largest cluster (green) comprised frontal and parietal areas (IPS1-4, SPL, FEF, and IFS) associated with the dorsal attention network (21). Within this cluster, the peaks of pulvino-cortical connectivity maps were topographically organized and reflected the spatial organization of cortical areas. Posterior parietal (IPS1/2) peaks were located most posteriorly in the dorsal pulvinar, followed by anterior parietal (IPS3/4) and then frontal regions (FEF and IFS). The peak for SPL, which is located medial to the IPS maps in cortex, was located medial to the IPS peaks in the pulvinar. The distances between the peaks in this cluster were correlated with the cortical distances between areas (r(19)=0.82, p<0.0001). A second cluster (red) contained IPS5 and regions in anterior parietal (antPPC) and lateral temporal (latTemp) cortex that we, and others, have shown to form a human-specific tool network(Chao and Martin, 2000; Mahon et al., 2007; Mruczek et al., 2013; Kastner et al., 2017). Within this cluster, there was no clear relation between the distances of pulvino-cortical connectivity peaks and cortical distance in this cluster. The third cluster (blue) contained two regions in the inferior parietal cortex (IPL) and the posterior cingulate / precuneus (PreC) that are associated with the default mode network(Buckner et al., 2008) as well as two additional regions in medial (medPPC) and lateral parietal cortex (latPPC) (See Materials and Methods: additional areas). Within this cluster, distances between peaks of pulvino-cortical connectivity reflected functional similarity, not cortical distance. The pulvino-cortical connectivity peaks of the functionally similar inferior parietal and posterior cingulate/precuneus areas were within 1mm of each other, and the other two lateral and medial parietal areas were within 0.5mm of each other. The peak of the lateral (and medial) parietal area and the peak of inferior parietal (and posterior cingulate/precuneus) area connectivity maps were both separated by several millimeters even though these areas were cortically adjacent. Further, the axes of organization within each cluster did not map across other clusters, suggesting that the organization of areas within a cluster did not translate to other clusters. For example, the anterior-posterior axis reflected cortical distance within the first cluster containing most IPS and frontal maps, but did not for the other two clusters. Last, it is notable that lateral and medial parietal areas, typically not considered to be part of visual cortex, were clustered together and distinct from other visual areas, suggesting that within the dorsal pulvinar there is a distinction in connectivity between visual and non-visual cortical regions. Taken together, these data suggest that the organization of connections between cortex and the dorsal pulvinar mainly reflects functional specialization of cortex and that multiple functionally dissociable sub-regions likely exist within the dorsal pulvinar.

**Figure 6.**
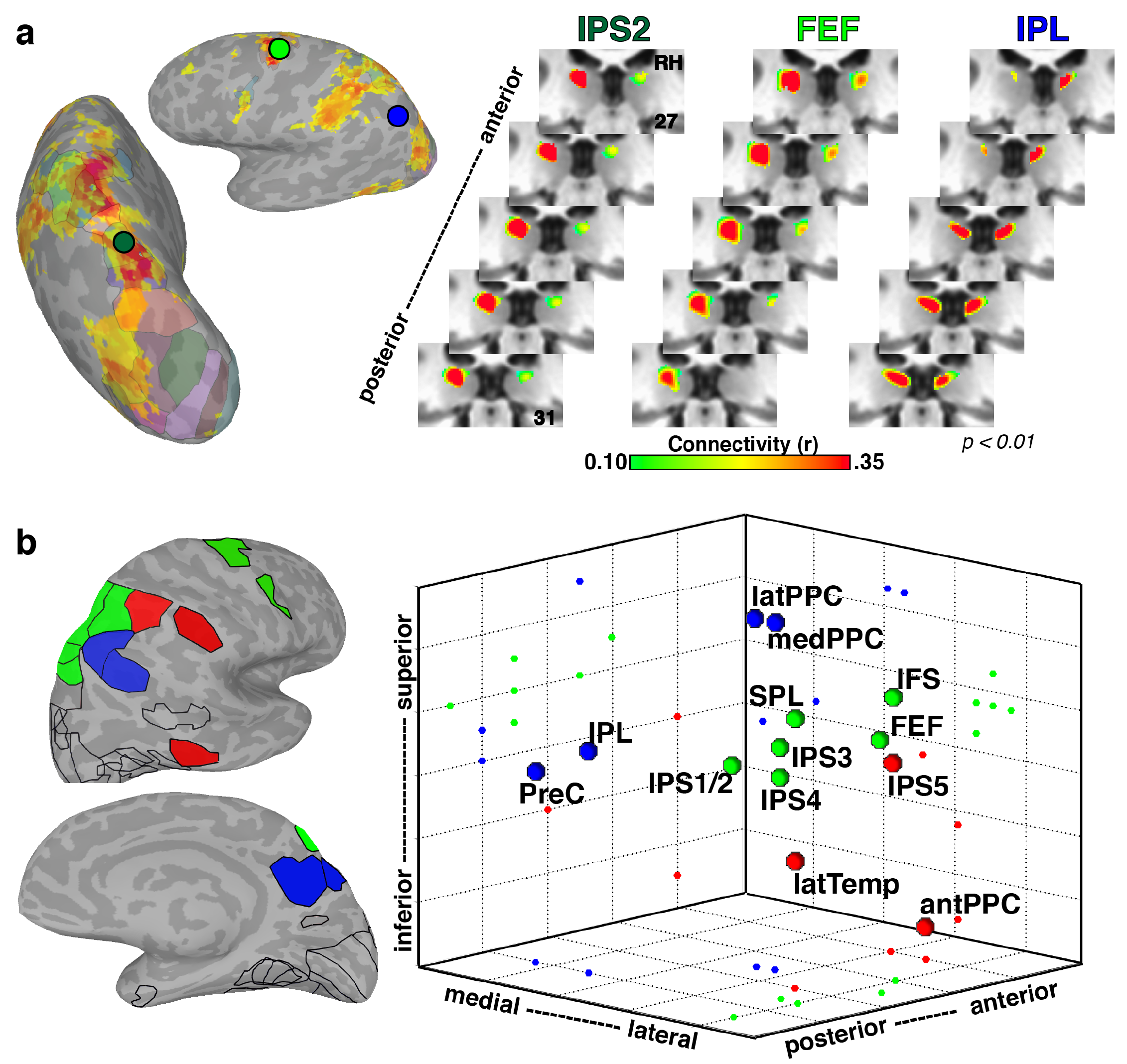
Functional connectivity with dorsal pulvinar reflects both functional organization and distance between cortical areas. (a) Group average (n=13; p < 0.01 from one-sample t-test across subjects) correlations within the dorsal pulvinar for two cortical seed areas (IPS2 and FEF as well as their overlap (yellow). Surface figures illustrate the cortical correlation patterns from IPS2 (blue) and FEF (red) seeds in a single subject. (b) Three-dimensional plot of cortical connectivity map peaks in the dorsal pulvinar. Peak coordinates from the left hemisphere were reflected across the midline and averaged with the right hemisphere. Pulvino-cortical connectivity within the dorsal pulvinar was clustered into three groups. Surface figures illustrate the outlines of all cortical areas tested and color filled areas correspond to those included in the clustering of the dorsal pulvinar. Within the largest cluster (green) containing the IPS maps, FEF, and IFS, connectivity reflected cortical distance with IPS1/2 located most posteriorly, followed by IPS3/4 and then FEF and IFS located anterior. The peak connectivity for SPL, which is located medial to the IPS maps cortically, was located medial to the IPS peak correlations in the pulvinar. A second cluster (red) contained tool-selective regions in anterior parietal and lateral temporal cortex as well as IPS5. The third cluster (blue) contained medial, lateral, and inferior parietal areas as well as the precuneus.

### Organization of cortical connections within the ventral pulvinar

Within the ventral pulvinar, the organization of pulvino-cortical connectivity broadly reflected both cortical distance and functional similarity. Consistent with prior work from our lab testing fewer areas (Arcaro et al., 2015), individual occipital and temporal areas were most strongly correlated with the ventral pulvinar (Fig. 4a). Functional connectivity of face-selective areas (OFA, FFA, and AT) and scene-selective area PPA was localized to the ventral posterior pulvinar (Fig. **7**a). In contrast to the dorsal pulvinar, overlap between pulvino-cortical connectivity maps was largest for cortical areas in close anatomical proximity (e.g., OFA and FFA) regardless of functional specialization (e.g., face-selective FFA and scene-selective PPA). Connectivity of OFA and FFA overlapped but was largely distinct from AT. Notably, the FFA’s functional connectivity was localized to the same postero-medial region of the ventral pulvinar that showed face-selective activity (Fig. 2d), demonstrating that pulvino-cortical connectivity patterns were predictive of the selectivity within the pulvinar. To evaluate the finer-scale structure of these maps, the spatial locations of the peaks in pulvino-cortical functional connectivity maps were assessed for each area. There was a general anterior-lateral to posterior-medial gradient in the location of pulvino-cortical connectivity peaks from posterior occipital to temporal cortical areas. Clustering of pulvino-cortical connectivity within the ventral pulvinar revealed 2 groups segmenting lateral and medial portions (Fig. 7b). This clustering grouped functionally similar areas. Face-selective areas FFA, OFA, and AT were grouped in the medial (blue) cluster and scene-selective areas PPA (also PHC1/2) and TOS were grouped in the lateral (red) cluster. However, the lateral and medial clusters also reflected anatomical distance, differentiating medial occipito-temporal cortex from lateral temporal cortex, respectively. Notably, the mapping of cortex along the medial-lateral axis was flipped such that medial cortical areas (e.g., PHC and VO) mapped onto the lateral pulvinar and lateral cortical areas (e.g., FFA and AT) mapped onto the medial pulvinar, which appears consistent with broad anatomical connectivity patterns between temporal cortex and the pulvinar in macaques(Benevento and Rezak, 1976). To further differentiate effects of functional specialization and cortical distance, a subset of areas within ventral temporal cortex was selected to directly compare face- and scene-selective regions as well as early (V1, V2, V3) and intermediate (hV4, VO1/2) visual cortex, which likely comprise the inputs. Clustering on this subset of areas yielded 4 clusters (Fig. 7c). One cluster (red) contained posterior occipital areas V1, V2, V3, and hV4. A second cluster (magenta) contained occipital-temporal areas VO1-2, PHC1, and PPA. A third cluster (blue) contained temporal areas FFA and AT and a fourth cluster (green) contained temporal area PHC2 and occipital-temporal area OFA. Though face-selective areas FFA and AT were clustered together, another face-selective area, OFA, was clustered with scene-selective area PHC2 and the distances between pulvino-cortical connectivity peaks were well explained by the cortical distances across all areas (r(64) = -0.72, p < 0.0001). These data suggest that in contrast to the dorsal pulvinar, pulvino-cortical connectivity with the ventral pulvinar predominantly reflected cortical distance.

Cortical distance relative to primary visual area V1 best accounted for the distances between pulvino-cortical connectivity peaks in the ventral pulvinar. To evaluate the influence of individual areas, instead of evaluating the relationship between cortical distance and pulvino-cortical connectivity peaks for all areas, comparisons were made relative to a single reference area. V1 as the reference area (i.e., the distances between each area and V1) yielded the strongest correlation between cortical distance and the peaks of functional connectivity (Fig. 7d, inset image). To illustrate this relationship, we plotted the distances of peak correlations for all occipital-temporal areas (Fig. 7d) and the subset of occipital, face-, and scene-selective areas (Fig. 7e) relative to V1. The distances for occipital-temporal areas fell close to a line between V1 and the maximally distant temporal area, AT. Taken together, these data suggest that cortical distance is a prominent organizing principle of cortical connections with the ventral pulvinar and the spatial layout of cortical connections appears to be anchored to primary visual area V1.

**Figure 7.**
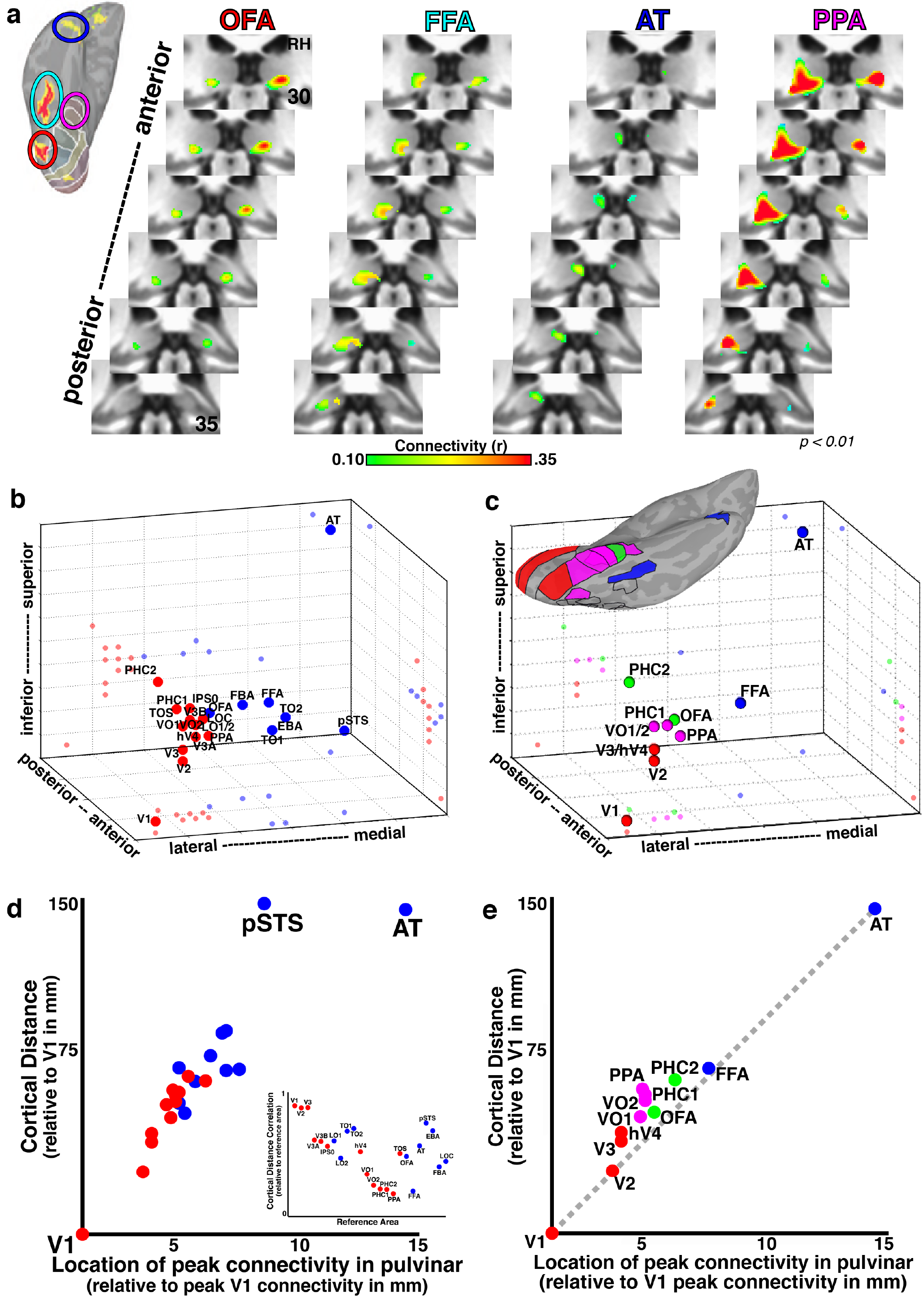
Functional connectivity with ventral temporal cortex was strongest within the posterior half of the ventral pulvinar. (a) Group average correlation maps (n=13; p < 0.01 from one-sample t-test across subjects) are shown for three cortical areas defined based on their functional specialization: occipital face area (orange), fusiform face area (green), and anterior temporal face area (purple). Surface figure illustrates the localization of face-selective regions in an individual subject. Each cortical region showed greater activity for face vs. place stimuli (p < .01, FDR-corrected). Correlations were strongest within the ventral and posterior-most portions of the pulvinar for each area. (b) Three-dimensional plot of the peak occipital and temporal pulvino-cortical connectivity within the ventral pulvinar. Spheres depict the 3D spatial location of each area’s peak connectivity within the ventral pulvinar. 2D projections of each data point are plotted on the walls and floor of the graph. Clustering revealed 2 groups that differentiated lateral (red) and medial (blue) portions of the ventral pulvinar. (c) Same plot as in (b), but for a subset of ventral areas. Clustering of connectivity within the ventral pulvinar revealed 4 groups. (d) A scatter plot of cortical distance relative to V1 vs. distance of peak location relative to V1 in the pulvinar for all 24 areas in (b). Inset graph shows the cortical distance correlation relative to each of the ventral areas. (e) Same plot as in (d), but for the subset of areas in (c).

## Discussion

The functional organization of the human pulvinar reflected several distinctions between the dorsal and ventral visual pathways in cortex. Similar to occipital and ventral extrastriate cortex, the ventral pulvinar responded preferentially to contralateral visual space. These results are consistent with previous work from our lab demonstrating that the human pulvinar contains at least two maps of visual space(Cotton and Smith, 2007; Arcaro et al., 2015; also see DeSimone et al., 2015), similar to other primate species(Allman et al., 1972; Gattass et al., 1978; Bender, 1981; Li et al., 2013). The ventral pulvinar also showed consistent responses to dynamic movie stimuli even when frames were presented out of order, disrupting the temporal structure and narrative(Hasson et al., 2008), suggesting that the ventral pulvinar tracks moment-to-moment variations in low-level visual features similar to early visual cortex. In contrast, the dorsal pulvinar was visually responsive to both contralateral and ipsilateral visual space, but showed increased activity during directed spatial attention similar to fronto-parietal cortex(Kastner et al., 1999; Szczepanski et al., 2010; also see Fischer and Whitney, 2012). The dorsal pulvinar only showed consistent responses to repeated presentations of intact movies, not to a temporally scrambled version of the movie, suggesting that it was generally insensitive to low-level stimulus features and likely is involved in higher-level cognitive processes such as tracking the narrative and attentional state. These functional differences between the dorsal and ventral pulvinar suggest that the visual functions of the pulvinar are quite varied, similar to cortex.

The ventral pulvinar responded to high-level visual information. A posterior medial region of the ventral pulvinar responded preferentially to face stimuli (vs. scenes) and was functionally coupled with the fusiform face area at rest. A lateral region of the ventral pulvinar showed weak selectivity for scenes (vs. faces) and was functionally coupled with scene-selective areas at rest. These data are consistent with anatomical connectivity studies in macaques that demonstrated projections from the medial ventral pulvinar to the lower lip of the STS(Grimaldi et al., 2016) and projections from ventral temporal cortex (around area TF) to lateral portions of the pulvinar(Benevento and Rezak, 1976) as well as an electrical stimulation study showing evoked activity within the ventral medial pulvinar from stimulation of face patches in the lower bank of the STS (Moeller et al., 2008). Previous studies have demonstrated responsiveness in the primate pulvinar to object features at the neuronal level (Nguyen et al., 2013; Nguyen et al., 2016). To our knowledge, this is the first demonstration of image category-selective visual responses within the human thalamus.

Functional distinctions between the dorsal and ventral pulvinar were further characterized based on their connections with visual cortex. The pulvinar was correlated with 39 functionally dissociable areas across occipital, parietal and temporal cortex. Individual cortical areas were correlated with focal regions of the pulvinar. Correlations of neighboring cortical areas tended to overlap in the pulvinar. Individual areas were functionally coupled to either the dorsal or ventral pulvinar, but generally not both. The ventral pulvinar was functionally coupled with occipital and temporal cortex at rest, consistent with anatomical studies in other primate species (Baizer et al., 1993). The dorsal pulvinar was functionally coupled with parietal and frontal cortex, consistent with anatomical studies in other primates (Yeterian and Pandya, 1985; Schmahmann and Pandya, 1990; Hardy and Lynch, 1992). These results are consistent with a previous study where we only considered connectivity with a subset of regions and did not assess the relationship of pulvino-cortical connectivity patterns between cortical areas (Arcaro et al. 2015). Our correlation results bridge the distribution of functional response properties across the human pulvinar that we report here with the organization of pulvino-cortical anatomical connectivity previously reported in monkeys. Our results suggest that the human pulvinar and its communication with visual cortex is largely segregated into two pathways similar to other primates (Figs. 2 & S3; Baleydier and Morel, 1992; Baizer et al., 1993; Kaas and Lyon, 2007), and emphasizes that this important organizing principle of the visual system is not restricted to cortex.

The spatial organization of pulvino-cortical interactions reflected two principles: cortical distance and functional similarity. In general, sub-regions of the pulvinar were correlated with cortical regions that have similar response properties. Within the ventral pulvinar, correlation patterns predominantly reflected distances between cortical areas. Consistent with prior anatomical studies in monkeys (Webster et al., 1993; Adams et al., 2000; Shipp, 2003), correlation peaks for occipital and temporal seed areas were distributed along a rostrolateral to caudomedial axis with V1 correlations located rostral and anterior temporal area, AT, located caudal. There was a near perfect linear relationship between the distance of each area’s peak functional connectivity in the pulvinar and cortical distance relative to V1. While some aspects of the cortical correlation patterns within the ventral pulvinar were consistent with the functional specializations of occipito-temporal cortical areas, these were generally also accounted for by anatomical distance. For example, correlations of two face-selective areas partially overlapped within the ventral pulvinar and were distinct from correlations with a scene-selective area, though these data reflected broader differences between ventral-lateral and ventral-medial cortex. The lack of clear clustering of pulvino-cortical correlations within the ventral pulvinar based on functional specialization may appear at odds with our finding of image category selective clusters within the ventral pulvinar, which suggests the presence of some functional clustering. However, this likely reflects a difference of spatial scale (local vs. interareal). Local clustering of functionally similar neurons (e.g., a face patch) reflects a minimization of cortical distance. Therefore, functional clustering of response properties within the ventral pulvinar (e.g., face-selective cells) is not at odds with the cortical distance. This is further supported by our findings that a face-selective region of the ventral pulvinar was correlated with a single, functionally similar cortical region (e.g., FFA) while correlations with anatomically separated, but functionally similar area AT did not converge. In contrast to the ventral pulvinar, correlations between the dorsal pulvinar and fronto-parietal cortex predominantly reflected functional similarity. Spatial clustering of fronto-parietal cortical correlations within the pulvinar differentiated the dorsal attention network from the default mode and anterior parietal/tool networks. Further, pulvinar correlation maps for areas such as IPS2 and FEF largely overlapped and were clustered despite their large cortical distance. At a finer scale, cortical distance was apparent in the spatial organization of correlations between the dorsal pulvinar and a subset of fronto-parietal cortical areas. Within the dorsal attention network cluster, the peaks of cortical functional connectivity maps within the pulvinar were roughly distributed along an anterior-posterior axis similar to relative cortical distances. However, this relationship was not observed for the other two clusters within the dorsal pulvinar. Altogether, these results suggest that the pulvinar’s influence on cortex varies between its dorsal and ventral subdivisions.

The pulvinar has been shown to regulate communication within and between cortical areas (Purushothaman et al., 2012; Saalmann et al., 2012). We propose that the dorsal pulvinar predominantly facilitates communication between widespread cortical areas that mediate top-down processes such as control of attention, as well as the default mode and temporo-parietal tool networks, and the ventral pulvinar predominantly facilitates cortically local communication between areas involved in visual recognition and feature extraction. Similar to cortex, interactions between dorsal and ventral pulvinar likely exist, though these were not prominent in our data with the exception of retrosplenial area RSC, which appeared to be a hub linking the pulvinar subdivisions. Overall, our results suggest that the human pulvinar represents the entire visual cortex and the principles that govern its organization, though in a compressed form.

## ACKNOWLEDGMENTS

This work was supported by grants from the National Institutes of Health (RO1 MH064043, RO1EY-017699, and T90DA-022763) and the National Science Foundation (BCS-1328270). We thank Christopher Honey, Stephanie McMains, and Ryan Mruczek for help with data collection.

